# Age-related seroprevalence trajectories of seasonal coronaviruses in children

**DOI:** 10.1101/2022.07.26.501649

**Authors:** Yasha Luo, Huibin Lv, Shilin Zhao, Yuanxin Sun, Chengyi Liu, Chunke Chen, Weiwen Liang, Kin-on Kwok, Qi Wen Teo, Ray TY So, Yihan Lin, Yuhong Deng, Biyun Li, Zixi Dai, Jie Zhu, Dengwei Zhang, Julia Fernando, Nicholas C Wu, Hein M. Tun, Roberto Bruzzone, Chris KP Mok, Xiaoping Mu

**Affiliations:** Department of Clinical Laboratory, Guangdong Women and Children Hospital, Guangzhou, China; HKU-Pasteur Research Pole, School of Public Health, Li Ka Shing Faculty of Medicine, The University of Hong Kong, Hong Kong SAR, China; Li Ka Shing Institute of Health Sciences, Faculty of Medicine, The Chinese University of Hong Kong, Shatin, Hong Kong SAR, China; The Jockey Club School of Public Health and Primary Care, The Chinese University of Hong Kong, Hong Kong SAR, China; Carl R. Woese Institute for Genomic Biology, University of Illinois at Urbana-Champaign, Urbana, IL 61801, USA; Department of Chemistry and The Swire Institute of Marine Science, The University of Hong Kong, Pokfulam Road, Hong Kong, China; Department of Biochemistry, University of Illinois at Urbana-Champaign, Urbana, IL, 61801, USA; Center for Biophysics and Computational Biology, University of Illinois at Urbana-Champaign, Urbana, IL 61801, USA; Carle Illinois College of Medicine, University of Illinois at Urbana-Champaign, Urbana, IL 61801, USA

## Abstract

Four seasonal coronaviruses, including HCoV-NL63 and HCoV-229E, HCoV-OC43 and HCoV-HKU1 cause approximately 15–30% of common colds in adults. However, the frequency and timing of early infection with four seasonal coronaviruses in the infant are still not well studied. Here, we evaluated the serological response to four seasonal coronaviruses in 1886 children under 18-year-old to construct the viral infection rates. The antibody levels were also determined from the plasma samples of 485 pairs postpartum women and their newborn babies. This passive immunity waned at one year after birth and the resurgence of the IgGs were found thereafter with the increase of the age. Taken together, our results show the age-related seroprevalence trajectories of seasonal coronaviruses in children and provide useful information for deciding vaccine strategy for coronaviruses in the future.

## Main Text

SARS-CoV-2 is now high prevalence worldwide and becomes persistence in the human population. It is reasonable to expect that most people will be exposed to the virus for the first time during their childhood. Understanding the development of acquired immunity against the seasonal coronaviruses (HCoV-NL63 and HCoV-229E, HCoV-OC43 and HCoV-HKU1) in young age group will thus give us a clue on the impact of SARS-CoV-2 to human in the post COVID-19 era. These viruses have been circulating in human population for many years and are accounted for approximately 15–30% of upper respiratory tract infection^1^. Infection of these viruses mainly cause self-limiting flu-like illnesses, but severe pediatric respiratory infections are not rare^2-4^.

Children are not entirely immunological naïve when they are born^5^. IgG antibodies in the neonates are transferred from their mother so as to provide a transient immune barrier against the potential infection^6,7^. This transferred immunity plays a protective role before the infants establish their own specific adaptive immunity to the same pathogen. So far, there is paucity of data to describe the transition period from transferred immunity to acquired immunity for the seasonal coronavirus in children. Moreover, the accumulation of immune response to the seasonal coronaviruses in children is also not yet well understood. Longitudinal study showed that adults are repeatedly infected by the seasonal human coronaviruses for every 12 months^8^. Although it was found that the induction of antibodies after each infection is short-lasting, frequent reinfections lead to persistent levels of antibodies to the four seasonal coronaviruses in most of the adults^9^. These pre-existing antibodies against seasonal coronaviruses were recently found to be associated with the neutralizing antibody response against SARS-CoV-2 that may mitigate disease manifestations from SARS-CoV-2 infection^10^. In this study, we determined the serological response against four seasonal coronaviruses in the plasma samples of children and modelled the seroprevalence trajectories of the four virus subtypes during the whole childhood period.

We tested the seroprevalence to the four seasonal coronaviruses by the Enzyme-linked Immunosorbent Assay (ELISA) using the plasma samples collected from 1886 children (Female: 43.9%) with age ranging from 0 (Neonates) to 18 years old in Guangzhou, China between January and March in 2020. Among our cohort, 259 were under 6 months old, 161 were 6 months-1 year old, 278 were 1-2 years old, 603 were 3-6 years old, 466 were 7-12 years old, 119 were 13-18 years old (Supplementary Table 1). The spike (S) protein of coronavirus, which plays an essential role in the receptor recognition and cell membrane fusion process, is composed of two subunits, S1 and the stalk-like S2^11^. Since there are 63– 98% of sequence similarity in the S2 among the seven human coronaviruses^12,13^, we specifically targeted to detect the level of IgG antibody to the S1 (HCoV-229E, HCoV-NL63, HCoV-HKU1) or hemagglutinin-esterase (HE) (HCoV-OC43) of the viruses.

The IgG levels to the four seasonal coronaviruses were determined from each plasma sample. The association between the IgG level and the age in each seasonal coronavirus was constructed by generalized additive models (GAM) (Figure 1)^14^. The restricted cubic splines (smooth curve) with five knots were used to visualize the association. We found that the seroprevalences of the four seasonal coronaviruses showed a similar trajectory. Compared to the entire childhood period, the levels of IgG in the neonates dropped significantly and reached to the lowest level after the age of 1 year (1.25 years: HCoV-229E; 1 years: HCoV-OC43; 1.08 years: HCoV-NL63; 1.08 years: HCoV-HKU1). The levels of IgG were then increased and accumulated when the children became older in age. The IgG levels against HCoV-OC43, HCoV-NL63 and HCoV-HKU1 were increased to the comparable levels of the neonates at the age of 8, 9 and 6 years respectively. However, it was intriguing to find that the IgG to the HCoV-229E was increased slower than the other seasonal coronaviruses and it reached to the comparable level of the neonates at the age of 16 years. Thus, our results implicate that the frequency of repeated infection of HCoV-229E was lower than that of the other three subtypes^15^. The serological results of each coronavirus were further stratified into two sex groups (male/female) and were further compared (Supplementary Figure 1). Importantly, we found that the IgG waning of all four seasonal coronaviruses in male neonates were much faster than that in female. The time required for dropping the IgG of each coronavirus to their lowest level in male neonates were 1.89 (HCoV-229E: 0.75(M) vs 1.42(F)), 1.89 (HCoV-OC43: 0.62(M) vs 1.17(F)), 1.72 (HCoV-NL63: 0.68(M) vs 1.17(F)), 1.75 (HCoV-HKU1: 0.67(M) vs 1.17(F)) folds faster than that of the female neonates.

**Figure 1.**
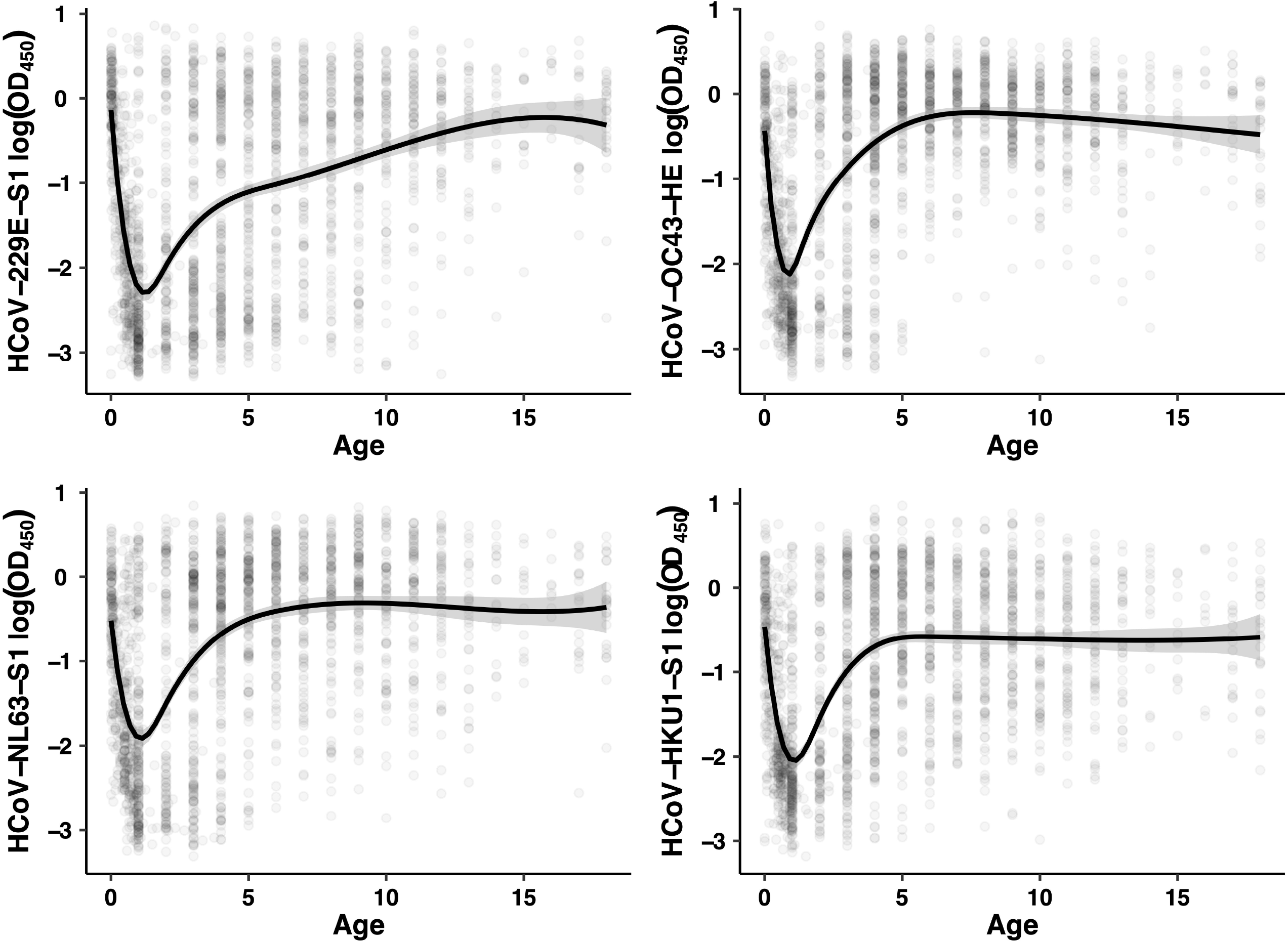
Seroprevalence trajectory of the four seasonal coronaviruses in children. The plasma samples were collected from 1886 children who aged from 0 (neonates) to 18 years old. Each sample was tested by ELISA against either S1 (HCoV-229E, HCoV-NL63 or HCoV-HKU1) or hemagglutinin-esterase (HCoV-OC43) protein. Generalized addictive models (GAM) was used to model the association between the serological data and the age. The black lines showed the fitted values and gray areas showed the 95% confidence intervals. Each sample was tested in duplicate, and the results were represented by the mean of the two values.

The relatively high levels of IgG antibody to the four seasonal coronaviruses in the neonates implicated a vertical transfer of the maternal immune response. It has been recently shown that the passive immunity against SARS-CoV-2 of neonates was contributed by their mothers^6^. We further collected plasma samples from 485 pairs of postpartum women and their newborn baby for testing the levels of their IgG to the four seasonal coronaviruses using similar serological assays. We found that the maternal IgG level was linearly associated with their neonatal IgG levels in each seasonal coronavirus: HCoV-229E (r=0.63, 95% CI: 0.57-0.68, *p*<0.0001), HCoV-OC43 (r=0.65, 95% CI: 0.60-0.70, *p*<0.0001), HCoV-NL63 (r=0.69, 95% CI: 0.64-0.74, *p*<0.0001), HCoV-HKU1 (r=0.63, 95% CI: 0.58-0.69, *p*<0.0001) (Figure 2). While comparing to the previous report that maternally derived antibodies against SARS-CoV-2 could persist up to 6 months of age in their infant^16^, our results indicated that the passive transferred immunity against the seasonal coronaviruses in neonates can maintain longer time.

**Figure 2.**
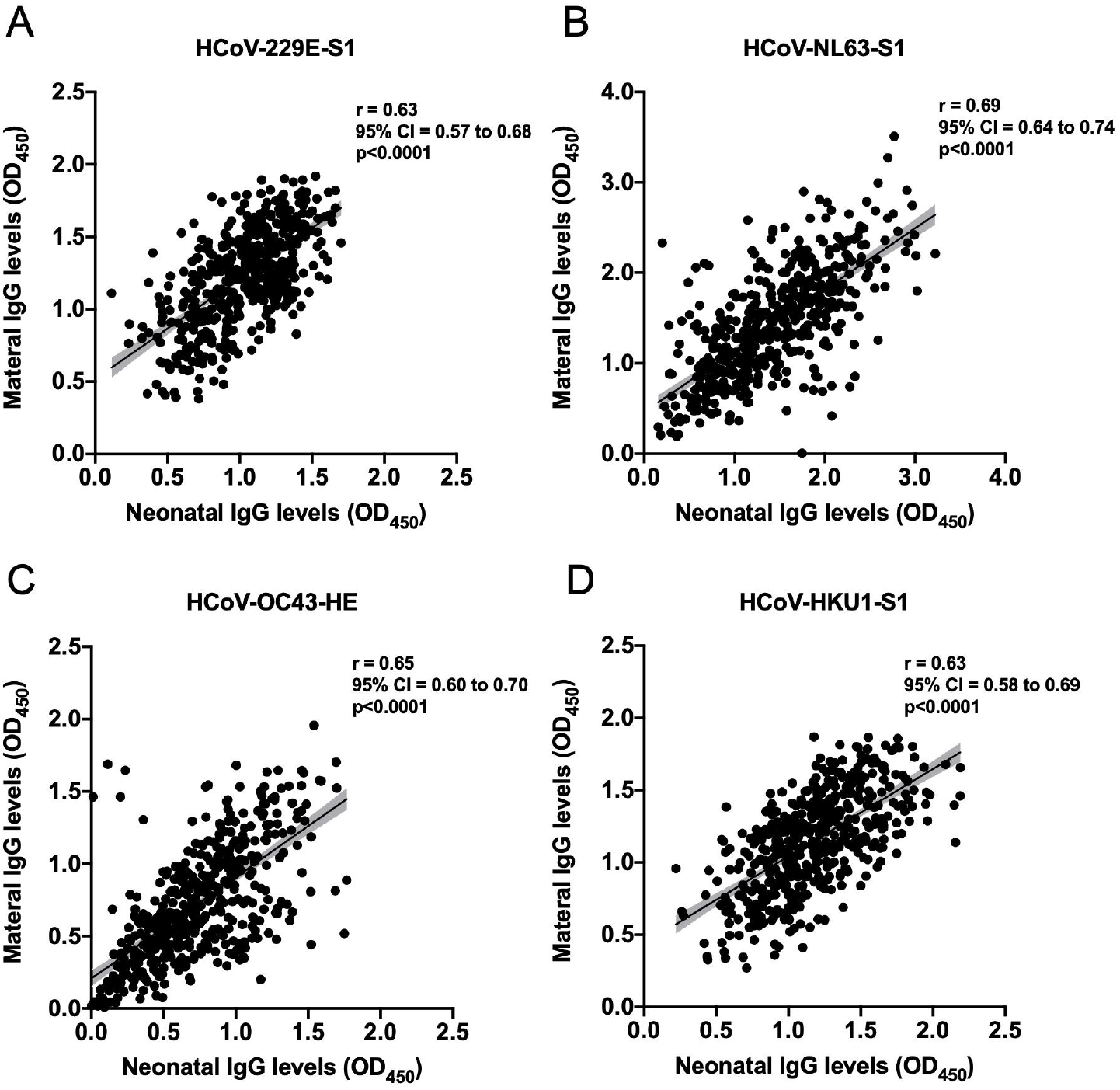
Correlation between the maternal and neonatal IgG levels of the four seasonal coronaviruses. 485 paired of maternal and neonatal plasma samples were collected and tested by ELISA. Antibody levels against A) HCoV-229E-S1, B) HCoV-NL63-S1, C) HCoV-OC43-HE, and D) HCoV-HKU1-S1 were determined and the correlations between the paired samples in the four seasonal coronavirus groups were shown. The black lines showed the fitted values and gray areas showed the 95% confidence intervals. The r represented the correlation coefficient.

Prevalence of the seasonal coronaviruses in children is determined either by detecting the specific nucleic acids from the respiratory specimen or through serology test. However, it is difficult to define and collect true negative reference samples because the seasonal coronaviruses are highly circulating in children. Previous studies adopted an approach in which the cutoffs were determined from a small subset of reference samples who the children were between 1-2 years old, and the tested samples were defined as positive if the results were above the mean of the references^17,18^. Here, we estimated the prevalence of the seasonal coronaviruses by using the lowest level in the generalized additive models as our negative reference (Supplementary Table 1). We assumed that children with IgG level above this point indicate infection of the corresponding seasonal coronaviruses and thus defined it as seropositive. 91.12%, 82.24%, 79.92% and 84.17% of sero-positivity to HCoV-229E, HCoV-OC43, HCoV-NL63 and HCoV-HKU1 respectively were found in those under 6 months old. In infants with the age between 6 months and 1-year-old, the seropositive rates dropped to 44.72% (HCoV-229E), 43.48% (HCoV-OC43), 45.96% (HCoV-NL63) and 45.96% (HCoV-HKU1). The sero-positivity of each seasonal coronavirus increased with age and was over 64.51% of prevalence in the children at their pre-school age (3-6 years). The seroprevalences for HCoV-229E, HCoV-OC43, HCoV-NL63 and HCoV-HKU1 kept increasing and were 98.11%, 100%, 96.23% and 98.11%, respectively, at the age of 16-18 years.

Our study described the transition from passive to acquired immunity for seasonal coronaviruses in children. The established approach here provides a view to identify the waning period of immunity against coronavirus after born, that will be useful to apply on SARS-CoV-2. The best timing to receive COVID-19 vaccine is still under debated. Though US CDC suggested that COVID-19 vaccination is recommended for children aged 6 months or older, it is mainly based on the safety concern rather than aiming for better protection. Defining the waning period in SARS-CoV-2 using our approach will provide scientific evidence to determine the vaccination window for children in post-COVID era. There were some limitations in our study. Firstly, the trajectories were illustrated using cross-sessional samples from population age groups, not in the longitudinal cohort. Secondly, the seroprevalences from our cohort were determined by ELISA only. The neutralizing effect to the seasonal coronaviruses was not evaluated. Thirdly, although the children were recruited from the non-respiratory ward or routine body check center, we did not collect their clinical background for analysis in this study.

In conclusion, we described that IgG antibody against four seasonal coronaviruses could be transferred from mother to their infant in a large-scale cohort. Importantly, we reported this transferred immunity waned for one year after born and children could acquire immunity against four seasonal coronaviruses with the increase of the age. Overall, these results provide a comprehensive analysis of the antibody dynamic in the early life of the children.

## Methods

### Sample Collection

Pediatric patients in non-respiratory diseases wards and children aged under 18 and without signs of influenza-like illness were recruited in our study. All plasma samples were obtained from the EDTA anti-coagulated peripheral blood samples in the Guangdong Women and Children Hospital, Guangzhou, China. Peripheral whole blood samples were centrifuged at 3000 x g for 10 minutes at room temperature for plasma collection. All plasma samples were kept in -80°C until used. Moreover, 500 plasma samples from postpartum women were collected between January and March 2020, with paired plasma samples collected from their newborn babies. All study procedures were performed after informed consent. The study was approved by the Human Research Ethics Committee at the Guangdong Women and Children Hospital (Approval number: 202101231).

### ELISA

The S1 subunits of spike protein (His tag) of HCoV-229E, HCoV-HKU1, and HCoV-NL63 and the hemagglutinin esterase protein (His Tag) of HCoV-OC43 were purchased from Sino Biological (China). The experiments were carried out according to our previous study^17^.

### Modelling

Generalized addictive models (GAM) was fitted to investigate the association between age and the ELISA results. The restricted cubic splines (smooth curve) with five knots were used to construct the model^14^. Of note, percentile places knots at five spaced percentiles of the explanatory variable, which are the 5th, 27.5th, 50th, 72.5th and 95th percentile. R version 4.0.4 was used for the analysis.

### Statistical Analysis

Significance between two groups was determined by the Mann-Whitney test, with a *p*-value smaller than 0.05 being considered statistically significant. Correlation between plasma samples were evaluated by using Pearson’s correlation coefficients.

## Acknowledgements

This work was supported by Calmette and Yersin scholarship from the Pasteur International Network Association (H.L.), Guangdong-Hong Kong-Macau Joint Laboratory of Respiratory Infectious Disease (20191205) (C.K.P.M.) and Emergency Key Program of Guangzhou Laboratory (Grant No. EKPG22-30-6) (C.K.P.M.).

## Author contributions

H.L., N.C.W. and C.K.P.M. conceived the research idea and designed the study. Y.L., Y.S. C.L., Y.D., B.L. and X.M coordinated and carried out cohort recruitment. H.L., S.Z., K.K., C.K.P.M. and H.M.T., analyzed the data. Y.L., H.L., C.C., W.L, Q.W.T, R.T.Y.S., Y.L, Z.D., J.Z., D.Z. and J.F. performed the experiments. H.L., R.B., H.M.T., and C.K.P.M. wrote the manuscript.

---

**Supplementary table 1.**
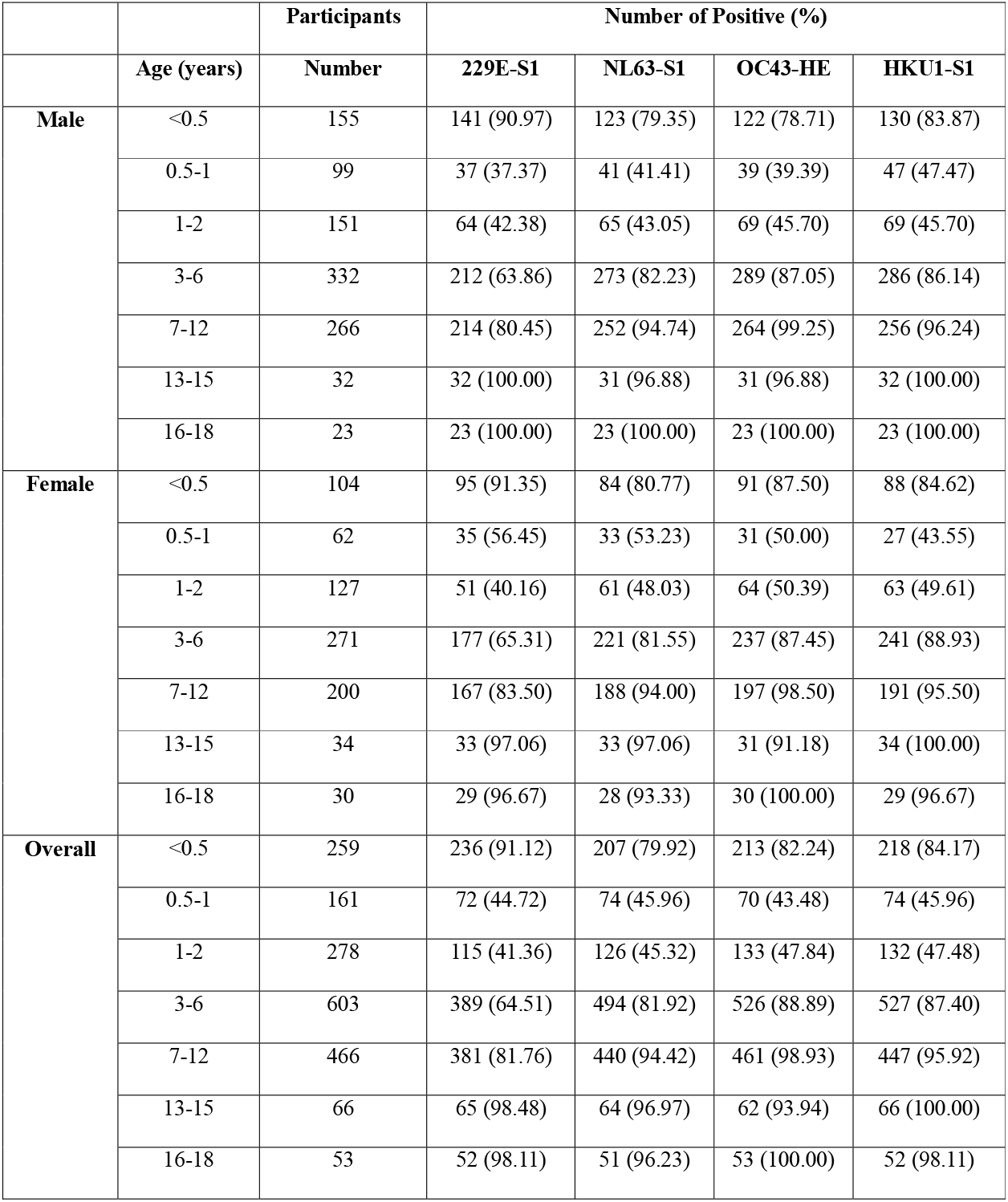
Prevalence of the seasonal coronaviruses in children.

## Figure legends

**Figure S1. Antibody levels against the four seasonal coronaviruses in different genders**. The 1886 plasma samples which were collected from children were further stratified into female (n=828 samples) and male (n=1058 samples) for analysis. Each sample was tested by ELISA against either S1 (A: HCoV-229E, C: HCoV-NL63 or D: HCoV-HKU1) or hemagglutinin-esterase (B: HCoV-OC43) protein. Generalized addictive models (GAM) was used to model the association between the serological data and the age. The black lines showed the fitted values and gray areas showed the 95% confidence intervals. Each sample was tested in duplicate, and the results were represented by the mean of the two values.

