## Supplementary figures and images for "Age-related seroprevalence trajectories of seasonal coronaviruses in children"

### Figure S1

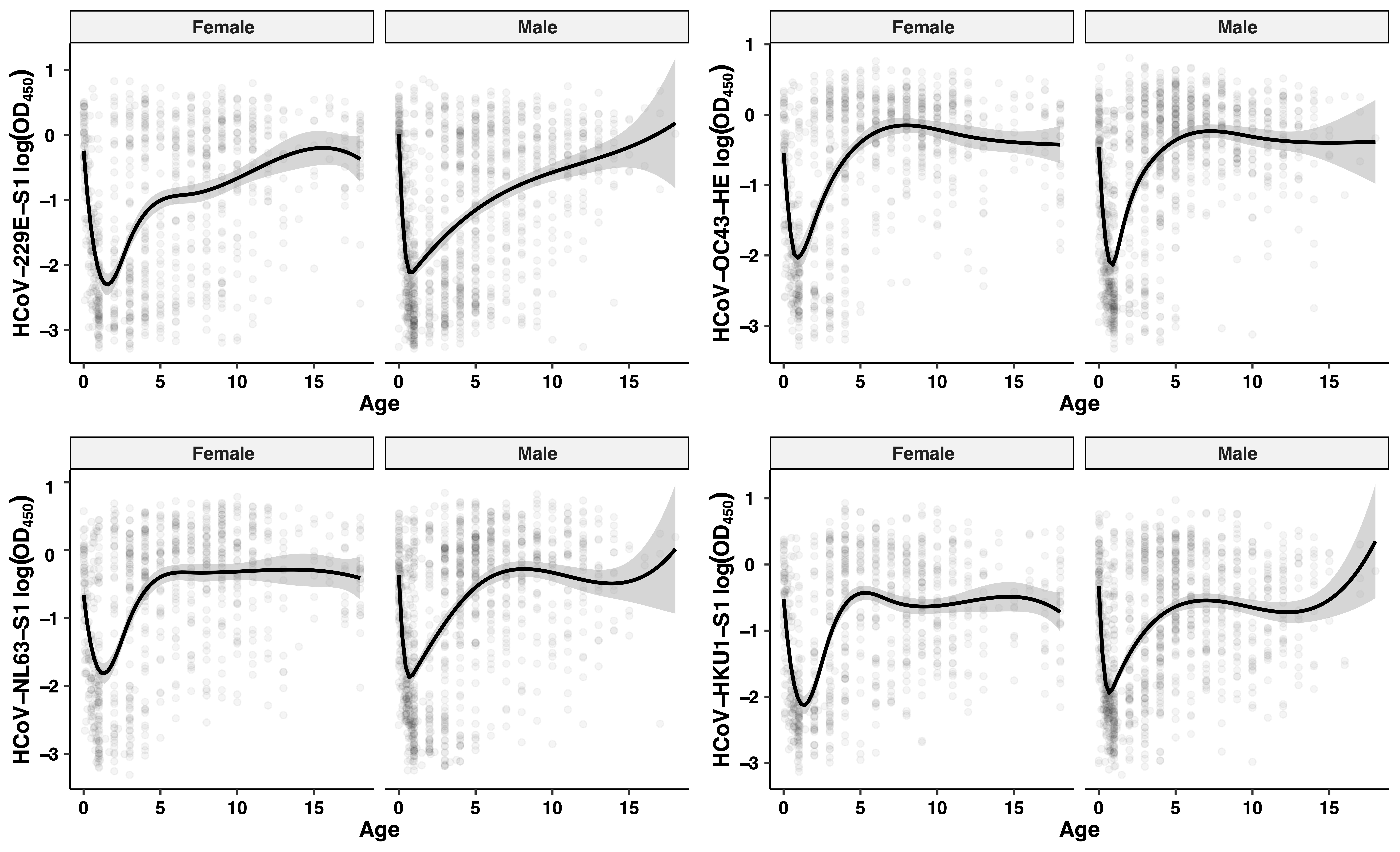
