## Supplementary material for "Age-related seroprevalence trajectories of seasonal coronaviruses in children": Table S1

**Supplementary table 1. Prevalence of the seasonal coronaviruses in children**

|  |  | **Participants** | **Number of Positive (%)** | | | |
| --- | --- | --- | --- | --- | --- | --- |
|  | **Age (years)** | **Number** | **229E-S1** | **NL63-S1** | **OC43-HE** | **HKU1-S1** |
| **Male** | <0.5 | 155 | 141 (90.97) | 123 (79.35) | 122 (78.71) | 130 (83.87) |
|  | 0.5-1 | 99 | 37 (37.37) | 41 (41.41) | 39 (39.39) | 47 (47.47) |
|  | 1-2 | 151 | 64 (42.38) | 65 (43.05) | 69 (45.70) | 69 (45.70) |
|  | 3-6 | 332 | 212 (63.86) | 273 (82.23) | 289 (87.05) | 286 (86.14) |
|  | 7-12 | 266 | 214 (80.45) | 252 (94.74) | 264 (99.25) | 256 (96.24) |
|  | 13-15 | 32 | 32 (100.00) | 31 (96.88) | 31 (96.88) | 32 (100.00) |
|  | 16-18 | 23 | 23 (100.00) | 23 (100.00) | 23 (100.00) | 23 (100.00) |
| **Female** | <0.5 | 104 | 95 (91.35) | 84 (80.77) | 91 (87.50) | 88 (84.62) |
|  | 0.5-1 | 62 | 35 (56.45) | 33 (53.23) | 31 (50.00) | 27 (43.55) |
|  | 1-2 | 127 | 51 (40.16) | 61 (48.03) | 64 (50.39) | 63 (49.61) |
|  | 3-6 | 271 | 177 (65.31) | 221 (81.55) | 237 (87.45) | 241 (88.93) |
|  | 7-12 | 200 | 167 (83.50) | 188 (94.00) | 197 (98.50) | 191 (95.50) |
|  | 13-15 | 34 | 33 (97.06) | 33 (97.06) | 31 (91.18) | 34 (100.00) |
|  | 16-18 | 30 | 29 (96.67) | 28 (93.33) | 30 (100.00) | 29 (96.67) |
| **Overall** | <0.5 | 259 | 236 (91.12) | 207 (79.92) | 213 (82.24) | 218 (84.17) |
|  | 0.5-1 | 161 | 72 (44.72) | 74 (45.96) | 70 (43.48) | 74 (45.96) |
|  | 1-2 | 278 | 115 (41.36) | 126 (45.32) | 133 (47.84) | 132 (47.48) |
|  | 3-6 | 603 | 389 (64.51) | 494 (81.92) | 526 (88.89) | 527 (87.40) |
|  | 7-12 | 466 | 381 (81.76) | 440 (94.42) | 461 (98.93) | 447 (95.92) |
|  | 13-15 | 66 | 65 (98.48) | 64 (96.97) | 62 (93.94) | 66 (100.00) |
|  | 16-18 | 53 | 52 (98.11) | 51 (96.23) | 53 (100.00) | 52 (98.11) |
